# Confocal-Compatible Workflow for Sectioning, Staining, and Imaging Serial Vibratome Sections for 3D Anatomical Reconstruction of the Lymph Node

**DOI:** 10.1101/2025.10.29.685350

**Authors:** Madeline A. Kibler, Margaux D. Miller, Michael J. Donzanti, Jason P. Gleghorn

## Abstract

3D anatomical reconstruction provides quantitative insight into tissue architecture, variability, and function. However, existing whole-organ imaging methods, such as light-sheet microscopy and micro-CT, are often costly, slow, and inaccessible for many laboratories. We present an accessible, confocal-compatible workflow for harvesting, sectioning, and staining murine lymph nodes for high-fidelity 3D reconstruction. The protocol preserves Z-plane resolution through vibratome sectioning and immunostaining, enabling detailed visualization of lymph node lobules and vasculature using standard confocal microscopes. The resulting image stacks can be used for quantitative morphometry and spatiotemporal pharmacokinetic modeling. This reproducible workflow expands access to spatial and structural studies of the lymph node and can be adapted to other small, heterogeneous organs.

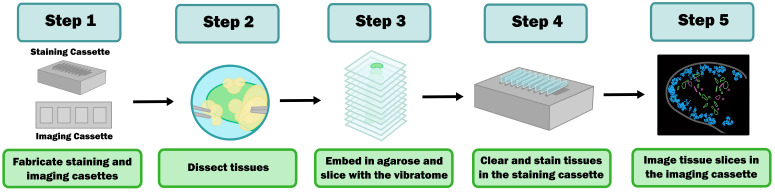

## Introduction

Anatomical reconstruction of whole organs and tissues is a growing area of interest in biological studies, as these 3D-reconstructed models can give accurate information on tissue architecture and organization, compartmentalization, and key physical interactions (1–5). 3D models can be used to quantify changes in geometry and morphology across developmental stages and disease states, variability within and between populations, and assess biological interactions with 3D spatial morphology (4, 6–10). Such reconstructions increasingly inform *in silico* modeling, drug distribution studies, and spatiotemporal analysis.

An organ with a strong need for anatomically accurate reconstruction methods is the lymph node (LN). LNs are small, heterogeneous, kidney bean-shaped organs located throughout the body, connected through a network of lymphatic vessels. LNs are highly heterogeneous and exhibit multiscale variability within and between individuals (11, 12). Architectural characteristics of a single LN can also change over time due to a variety of factors such as age and prior or ongoing infections (13). LNs maintain a highly compartmentalized architecture with specialized endothelial boundaries between compartments. These boundaries separate the circulating blood and lymph fluid from the parenchyma of the LN, including the cortex and paracortex, collectively called the lobule (14). This compartmentalization allows for a high rate of interaction between immune cells and antigens, significantly contributing to the LN’s role in adaptive immunity. However, it also complicates effective drug penetration to the LN, which is a growing area of research for LN-resident diseases such as metastatic cancers (15–24).

Here, we present a confocal-compatible, stepwise protocol for harvesting, sectioning, staining, and imaging murine lymph nodes for downstream 3D reconstruction. This approach provides an accessible and reproducible workflow that complements volumetric imaging systems, enabling high-fidelity anatomical reconstruction of the LN using standard confocal microscopes. The resulting 3D datasets can support spatiotemporal pharmacokinetic models, quantitative morphometry, and studies of immune architecture during development or disease.

### Comparisons to Other Methods

Various methods exist for 3D anatomical reconstruction, including complex imaging such as micro-computed tomography (µCT) and small-bore magnetic resonance imaging (MRI), serial histological sectioning, and whole-mount tissue imaging (2, 25–29). While these techniques are widely used in biological experimentation, many possess limited applicability for lymph node reconstruction. Small-bore MRI and µCT easily penetrate large tissues and provide high-fidelity data for reconstructions; however, a lack of contrast between soft tissues limits the ability to label multiple compartments within the lymph node without complicated machine learning techniques (30–33). Serial histological sectioning provides excellent X-Y resolution but is time-consuming, labor-intensive, and reagent-consuming to achieve comparable resolution in the Z-dimension, which is necessary for 3D reconstruction (34). To overcome this limitation, light-sheet microscopy and optical clearing approaches (e.g., CUBIC, vDISCO, Ce3D) have expanded the capacity for whole-organ imaging of immune tissues in 3D without sectioning or compromising Z-plane resolution (35). Yet these systems require expensive specialized optics and extended clearing protocols that limit accessibility (36, 37).

In contrast, the present protocol provides a complementary solution of a confocal-compatible, moderate-throughput workflow that preserves Z-plane resolution through vibratome sectioning and antibody staining. This allows 3D reconstruction using widely available imaging infrastructure and open-source analysis tools such as ImageJ, Imaris, or napari. This focus on accessibility and reproducibility distinguishes the workflow from prior volumetric approaches, making it particularly suited to broad adoption without advanced imaging infrastructure.

### Broader Applications

This protocol has been optimized for the visualization of the vasculature and lobule of the lymph node; however, because the workflow relies on confocal-compatible sectioning and standard immunostaining methods, it can be readily extended to other organs. Overall, this protocol provides an accessible approach to high-fidelity anatomical 3D reconstruction, applicable in understudied heterogeneous organs such as the uterus, placenta, pancreas, and tumors. In addition, this procedure can be paired with other applications, such as *ex vivo* tissue culture to quantify tissue structure after culture conditions or infection exposure, or to evaluate the effects of pharmacologic or cytokine treatment on microarchitecture (38–40). The modular design of the cassette and embedding workflow also enables adaptation for diverse antibody staining panels and downstream computational reconstruction using common laboratory resources. By emphasizing reproducibility and accessibility over instrumentation, this protocol complements light-sheet and whole-organ clearing methods, enabling access to spatial biology and anatomical mapping studies across laboratories equipped only with conventional confocal microscopes.

### Overview of the Procedure

This procedure was designed to produce high-fidelity 3D reconstructions of murine lymph nodes. As such, the following is a 5-step process: (1) fabrication of the tissue slice staining and imaging cassette, (2) harvesting the lymph node, (3) embedding and sectioning, (4) staining and optical clearing, and (5) imaging.

## Materials

### Cassette Fabrication

#### Reagents

- Isopropyl alcohol (Sigma-Aldrich, cat. no. W292907-1KG-K)
- Deionized (DI) water
- PDMS base (2065622, SYLGARDTM, Midland, MI)
- PDMS curing agent (2065622, SYLGARDTM, Midland, MI)
- Printing resin (TR250LV, Phrozen, cat. no. All_HIGHTEMP_RS)
- Resin Rinse (Form Wash, FormLabs, Somerville, MA)

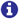 **NOTE:** Any 3D printer, monofilament or resin-based, should be able to generate the cassettes. The specific protocol enumerated herein is for resin-based printers with secondary curing and part rinsing/cleaning of uncured resin.

#### Equipment

- 100 mm Petri dish (Sigma Aldrich, cat. no. Z358762)
- Glass coverslip (VWR International, cat. no. 48393-106)
- 3D printer (Form 2, FormLabs, Somerville, MA)
- Resin Curing Oven (Form Cure, FormLabs, Somerville, MA)
- Alcohol Bath (Form Wash, FormLabs, Somerville, MA)
- Desiccator (5530000, Labconco, Kansas City, MO)
- Spin Coater (WS-650MZ-23NPPB, Laurell Technologies, Lansdale, PA)
- Plasma Cleaner (PDC-001-HP, Harrick Plasma, Ithaca, NY)

### Lymph Node Harvest

#### Biological Materials

- Small animal for LN harvest 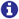 **NOTE:** This protocol has been validated with lymph nodes from wildtype and fluorescent reporter mice (Cdh5-Cre x mTmG, Jax Laboratories, #017968, #007676) previously. The procedure should be adaptable across organs from most small animal models and human tissues.

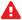 **CAUTION:** All animal studies and experiments using human tissues must comply with institutional and governmental regulations and guidelines with regard to informed consent and animal care. Animal studies herein were performed in accordance with Animal Use Protocols (AUP #1320 and #1367) approved by the Institutional Animal Care and Use Committee (IACUC) at the University of Delaware.

#### Reagents

- 70% ethanol in deionized water (Sigma-Aldrich, cat. no. 322415-100ML)
- 4% Paraformaldehyde in phosphate-buffered saline (PBS) (Thermo Scientific, cat. no. 322415-100ML)

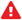 **CAUTION:** Paraformaldehyde is a carcinogenic, flammable, and a sensitizer. Take proper precautions to avoid inhalation and contact with skin or eyes.

- 0.1% Triton X-100 in PBS (Thermo Scientific, cat. no. AAA16046AE)
- 1X PBS (Corning, cat. no. 1482648)

#### Equipment

- *CO*_2_ chamber
- Forceps (Scientific Labwares, cat. no. PL-29)
- Dissection scissors (Sigma Aldrich, cat. no. S3146)
- Dissection board (Expanded Polystyrene)
- Dissection pins (Flinn Scientific, cat. no. AB1084)
- Stereoscope (Zeiss Axio Discovery V8, Oberkochen, Germany)
- Plastic wrap (Amazon Basics, cat. no. B09Z2LFBJC)
- Kimwipes® (KIMTECH, cat. no. B008N1YRZW)
- 5 mL macrocentrifuge tube (CELLTREAT, cat. no. 229449)

### Embedding and Sectioning

#### Reagents

- Low-melt agarose (IBI Scientific, cat. no. IB70051)
- 1X PBS (Corning, cat. no. 1482648)

#### Equipment

- 10×10×5 mm vinyl cryo-molds (TissueTek, Sakura, Torrance, CA)
- Vibratome (Campden 752M Vibroslice, Campden Instruments, Loughborough, United Kingdom)
- Razor blade (Feather, cat. no. B00301APUQ)
- Super glue (Loctite, cat. no. 2436365)
- Paint brush (Transon, cat. no. B0C6XQG5YL)
- Positive-displacement pipette
- 48-well plate (VWR International, cat. no. 10861-560)
- Scale (Precision Balance MS303TS/00, Mettler Toledo, Columbus, OH)
- Microwave

### Staining and Optical Clearing

#### Reagents

- 20X Phosphate-buffered saline (PBS) (Fisher Scientific, cat. no. 1482648)
- Double-distilled water
- Gelatine from bovine skin (Sigma-Aldrich, cat. no. G9391-100G)
- Bovine serum albumin (BSA) (Fisher BioReagents™, cat. no. BP9700100)
- Tween-20 (TCI Chemicals, cat. no. T0543-500G)
- Tris base (Sigma-Aldrich, cat. no. 93362)
- Sodium chloride (Sigma-Aldrich, cat. no. 71376)
- Hydrochloric acid (HCl) (Sigma-Aldrich, cat. no. H1758)

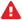 **CAUTION:** HCl is highly corrosive and can cause severe skin burns, eye damage, and respiratory irritation. Wear appropriate PPE and store in a corrosive-resistant container away from incompatible materials.

- Milli-Q® water or UltraPure water
- Sodium azide (Sigma-Aldrich, cat. no. S2002-5G)

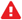 **CAUTION:** Sodium azide is highly toxic through skin, inhalation, and ingestion, and can cause eye, skin, and respiratory irritation. Use with proper ventilation. Take precautions when transferring powdered azide, as it is highly flammable and can easily dissipate with fume hood drafts, creating a dust cloud that can catch fire.

- Mouse Anti-lymphatic vessel endothelial hyaluronan receptor 1 (LYVE1) antibody (R&D Systems, cat. no. MAB2125-SP) Peripheral Node Addressin (MECA79) non-recombinant monoclonal antibody (Novus Biologicals, cat. no. NB100-77673SS)
- Platelet Endothelial Cell Adhesion Molecule-1 (CD31) antibody (Novus Biologicals, cat. no. NB600-562AF488)
- Donkey anti-Rat IgG AlexaFluor647 (Invitrogen, cat. no. A78947)
- Donkey anti-Rabbit IgG AlexaFluor488 (Invitrogen, cat. No. A21206)
- Donkey anti-Mouse IgG AlexaFluor594 (Invitrogen, cat. no. A21203)
- Tissue Clearing agent (Focus Clear, CelExplorer, Hsinchu City, Taiwan, cat. no. FC-101)

#### Equipment

- Glass beaker (United Scientific Supplies, cat. no. BG1000-500)
- 0.45 µm filtration bottle (VWR International, cat. no. 10040-438)
- Vacuum line or aspirator
- Glass coverslip (VWR International, cat. no. 48393-106)
- Aluminum foil (Amazon Basics, cat. no. B093X4M4QF)
- Confocal microscope (LSM800, Zeiss, EC PlnN 5x/0.16 Ph1 DIC0, Pln Apo 10x/0.45 Ph1 DICII and LD PlnN 20x/0.4 Ph2 DICII)
- Pipettes (Thermo Scientific™, cat. no. 14-386-312)
- Hot water bath
- Ice and ice bucket

### Confocal Imaging

#### Reagents

- Tissue Clearing agent (Focus Clear, CelExplorer, Hsinchu City, Taiwan, cat. no. FC-101)

#### Equipment

- Confocal microscope (LSM800, Zeiss, EC PlnN 5x/0.16 Ph1 DIC0, Pln Apo 10x/0.45 Ph1 DICII and LD PlnN 20x/0.4 Ph2 DICII)

### Reagent Prep

#### Blocking Buffer

##### Materials

- 20X PBS
- Double-distilled water
- Gelatine from bovine skin
- BSA
- Tween-20
- Glass beaker
- Scale
- Hot water bath or heated stir plate with stir bar
- 100-1000 µL pipette (P1000) and appropriately sized tips
- Scissors
- 0.45 µm filtration bottle
- Aspirator or vacuum line

##### Solution Preparation

1. Make 500 mL 1X PBS from 20X stock in clean beaker or glass bottle (25 mL 20X PBS + 475 mL double-distilled water).
2. Weigh 1 g of gelatine (for 0.1% solution) and add to 500 mL 1X PBS.
3. Place the glass bottle in a water bath or on a heated stir plate for 1 hour to dissolve gelatin.
4. Before adding BSA, let the solution cool to room temperature.
5. Weigh 5 g BSA (for a 0.5% solution) and add to the beaker while stirring.
6. Let BSA dissolve.
7. Cut off the tip of P1000 to pipette the viscous Tween-20.
8. Add 500 µL Tween-20 to the solution (final 0.1% solution).

##### Filtration

1. After all components are fully dissolved with no visible pellet, set up a 0.45 µm filtration bottle.
2. Pour 500 mL of solution into the top chamber of the filtration unit.
3. Attach the aspirator/vacuum line to the hose connector of the filtration unit.
4. Turn on the vacuum line to pull the solution from the top to the bottom chamber.
5. After all of the solution has been filtered, turn off the vacuum, unscrew the top chamber, and discard. Close the bottom chamber with a sterile cap (comes in the filtration unit packaging).

##### Storage

1. The bottom chamber has now become the storage container. Label the side with all components the solution contains, as well as a note to “*add 0*.*1% sodium azide when opening under unsterile conditions*”. Store at 4°C for future use.

#### Tris-buffered saline with 0.1% Tween-20 (TBST)

##### Materials

- Tris base
- Sodium chloride (NaCl)
- Milli-Q® or UltraPure water
- Hydrochloric acid (HCl)
- Tween-20
- Glass bottles large enough for 1L
- Scale
- pH probe
- P1000 and appropriately sized tips
- Scissors

##### Solution Preparation

1. In a glass bottle, dissolve 24 g of Tris base and 88 g NaCl in 900 mL of UltraPure water.
2. Insert a pH probe into the solution, and add HCl until the pH of the solution reaches 7.6.
3. Add water until a final volume of 1 L is reached to create 1 L of TBS 10X stock solution.
4. In a second glass bottle, combine 100 mL of the stock solution with 900 mL of UltraPure water.
5. Cut off the tip of a P1000 to pipette the viscous Tween-20.
6. Add 1000 µL Tween-20 to the solution (final 0.1% solution).

##### Storage

1. Solutions can be stored at room temperature for up to 1 week or at 4°C for up to 1 month. For longer storage, aliquot and store at −20°C. 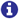 **NOTE:** The procedure above is for 1 L of stock solution and 1 L of TBST. For other volumes, refer to the calculators provided by Millipore Sigma (41).

### Procedure

#### 1. Cassette Fabrication

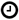 *Timing: Approximately 1-2 hours for fabrication and 18 hours for curing*.

##### Staining Cassette

1. Download the STL file of the negative mold of the staining cassette. See the Data and Resource Availability section at the end of the protocol for the URL.
2. Upload the STL file onto the Form Labs resin printer (**Figure 1A**).
3. Print the negative and remove it from supports with a blade before washing in isopropyl alcohol (Form Wash) and curing in the Resin Curing Oven (Form Cure).
4. Wash the negative in DI water and allow to air dry.
5. Mix polydimethylsiloxane (PDMS) at a base:curing agent ratio of 20:1.
6. Pour the PDMS over the mold in a shallow rectangular dish or petri dish. Ensure that the PDMS height does not fully submerge the mold, so that the mold can be removed after curing and tissue sections can be inserted into the completed cassette (**Figure 1B**). Degas the mixture by placing it in a desiccator and applying a vacuum until no bubbles remain. 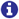 **NOTE:** Negative molds are designed to hold up to 5 tissue sections. Multiple molds may be aligned back-to-back to create staining devices with a number of positions that is a multiple of 5.
7. Bake overnight (18 hours) at 60°C.
8. Remove 3D-printed mold and cut around the PDMS form with dimensions less than a glass cover slip (**Figure 1C**). Be careful not to rip the fragile PDMS fins that divide the spaces for tissue slices when removing the mold (see Table **1**).
9. Mount the PDMS device to a glass microscope slide with plasma bonding. Place both the PDMS and cover slips bonding-side up in the plasma cleaner at 790-800 mTorr for at least 45 seconds. Immediately sandwich the cleaned surfaces together and apply light pressure to bond them together.

**Fig. 1.**
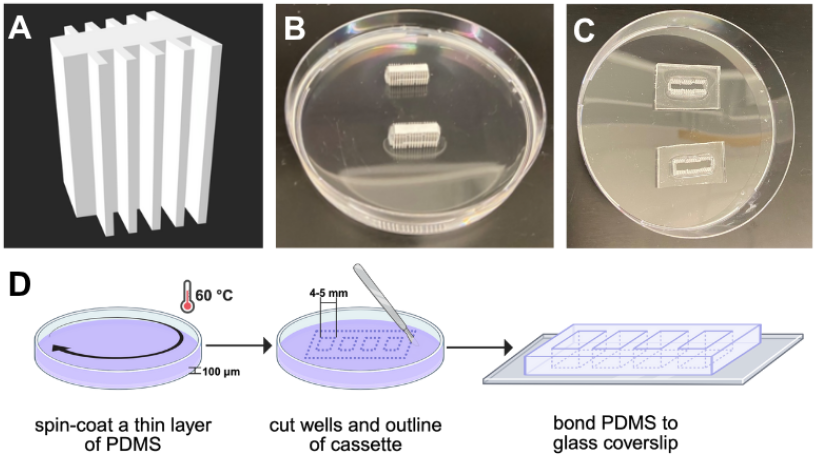
Fabrication of the staining and imaging cassettes. **(A)** 3D printed negative mold of staining cassette. **(B)** 3D printed molds are placed next to each other and embedded in PDMS to create 15 positions for tissue sections for each of the two staining cassettes. Once PDMS has cured, the **(C)** negative mold is removed and excess PDMS is trimmed away to create a domino-sized staining cassette. **(D)** To fabricate the imaging cassette, a thin layer of PDMS is spin-coated in a petri dish to define the PDMS thickness and allowed to cure before cutting four wells and bonding to a coverslip.

**Table 1.** Troubleshooting Table.

| Step | Problem | Possible Reason | Solution |
| --- | --- | --- | --- |
| 1 | PDMS fins rip during the removal of the negative mold. | Rough handling when removing mold. | Use sharp biological forceps to gently pull the PDMS off the mold. |
| 1 | No spin coater available. | Equipment not available. | Weigh out approximately 2.5g of PDMS in a 10 cm dish and allow it to fully spread across the surface before curing. This will produce approximately a 100 μm-thick PDMS layer for fabrication of the imaging cassette. |
| 2 | Difficulty finding lymph nodes. | LN is obscured by other tissues or an abundance of fat. | Grab a large section of tissue in the area where the LN is expected. Conservative rough dissection yields better results, followed by a fine dissection step. |
| 2 | Difficulty in fine dissection of lymph nodes. | Fat has dissolved into the PBS, obscuring the tissue and the target fat due to increased temperature and contamination of tissue from other LNs. | Move the organ into a fresh dish of ice-cold PBS. |
| 3 | Tissue is separating from the agarose block during vibratome sectioning. | The wt% of the agarose may not match the stiffness of the tissue sample. | Increase or decrease the wt% of agarose to better match the tissue's stiffness. |
|  |  | The speed and amplitude of the vibratome might not be optimal for the sample. | Start by reducing the vibratome speed and amplitude to prevent tearing of delicate tissues. Higher amplitudes are better for denser tissues, and lower speeds are better suited for softer tissues. Adjust as necessary for the specific tissue type and agarose wt%. |
|  |  | Dull vibratome blade. | Use a fresh vibratome blade for each sample, and if sectioning many samples at once, replace the blade periodically. |
|  |  | The organ is still lubricated with fat droplets. | Ensure that during fine dissection of a tissue, all fat has been removed from the surface of the organ. |
|  |  | The tissue has not been sufficiently dried. | After placing the tissue in the vinyl cassette, thoroughly dab the organ with a Kimwipe to take away water droplets and enhance agarose embedding. |
| 4 | Difficulty moving sections to and from the staining cassette. | Sections are fragile, and the agar block has likely been trimmed too short relative to the tissue. | Ensure wells are prefilled prior to adding sections to avoid wrinkling or folding of the section when dry. Use one or more forceps or a paintbrush when transferring sections to and from the cassette to avoid ripping the tissue or agar. |
| 4, 5 | Poor labeling quality. | The incubation time was too short. | Increase the incubation period for primary and secondary antibodies from 48 hours. It is safe to experiment with longer incubation times, as they will not significantly affect labeling quality once the stains have fully penetrated the tissue. |
|  |  | Incomplete antibody penetration deep into the tissue. | Increase the Triton-X concentration in the tissue fixative to 0.2% to improve sample permeability.<br>Increase antibody concentration to increase the gradient and improve transport deeper into the tissue. |
|  |  |  | Increase rocking speed during incubation. This will increase convection in the staining device and aid in stain penetration. |
| 5 | Laser is not completely penetrating the tissue. | Laser power is not high enough for the deeper portion of the stack. | Optically section the tissue in two domains: 1. the bottom half with the initial laser power, and 2. the top half with roughly 150% power. |

##### Imaging Cassette

1. Use a Spin Coater to spin coat a 100 mm petri dish with 5 g of 1:20 PDMS at 300 RPM for 30 seconds to achieve a coat of 100 µm thickness (**Figure 1D**). 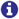 **NOTE:** Refer to Table **1** for how to achieve this thickness without the use of a Spin Coater.
2. Cure the PDMS overnight at 60°C.
3. Cut four *~* 4×5 mm rectangles in a row into the PDMS, then cut a large rectangle around the holes and peel off the resulting sheet from the petri dish. 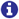 **NOTE:** The rectangular slots will be used to house lymph node slices during imaging, so you can adjust the sizing based on the tissue shape and size (**Figure 1D**).
4. Mount the PDMS to a glass coverslip by plasma bonding. Place both the PDMS and slides, bonding-side up, in the plasma cleaner at 790-800 mTorr for at least 45 seconds. Immediately sandwich the cleaned surfaces together and apply light pressure to bond them together.

#### 2. Lymph Node Harvest

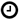 *Time:Approximately one hour*.

##### Gross Lymph Node Harvest

1. Prepare a dissection board by covering a piece of styrofoam in plastic wrap, fixing it in place with pins at the four corners of the board.
2. Prepare a Petri dish with 10 mL PBS. 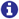 **NOTE:** Lymph nodes will be removed with large fat tissue borders and will be kept in PBS before fine dissection.
3. Euthanize the mouse in a *CO*_2_ chamber, followed by cervical dislocation.
4. Place mouse dorsal side down and pin the outstretched feet of the mouse down for maximum visibility of the ventral body cavity (**Figure 2A**).
5. Spray the mouse with 70% ethanol until the fur is saturated.
6. Lymph node harvest:
  a. Make an incision with scissors at the base of the abdomen, making a vertical incision to the top of the chest of the mouse. Maintain a shallow angle to avoid cutting anything but skin.
  b. Carry the incision distally to the left and right of the mouse, creating a Y-incision up the arms. Extend the incision down to the femur and the humerus of the mouse to expose the region behind the knee and the armpit.
  c. Cut connective tissue away from the body and the skin until the skin from the hip to the arm can be pinned down and away from the abdominal and chest cavities. (**Figure 2B**).
  d. Locate the LNs, and grip the surrounding fat with forceps, maintaining a wide margin, and cut out the lymph node, placing it in PBS. Repeat on the other side to harvest both left and right LNs. 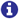 **NOTE:** Refer to **Figure 2A** for the location of the brachial, inguinal, and popliteal LN in a murine model. 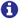 **NOTE:** Take care to not to cut any vessels in these regions, as it will obscure the dissection area. 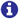 **NOTE:** The popliteal lymph node is small and not the most optimal in terms of size, cell count, and signal for downstream processing.

**Fig. 2.**
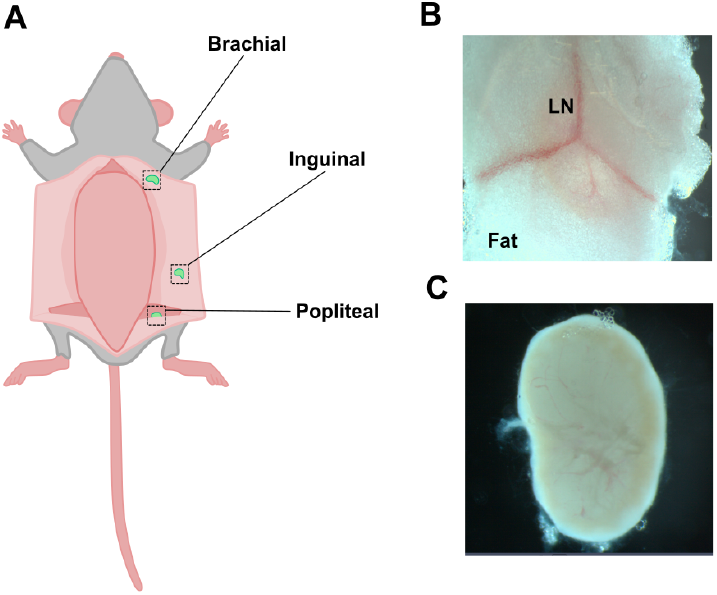
Lymph node dissection. **(A)** Location of target LNs in a mouse model. **(B)** Grossly dissected LN embedded in a fat pad. **(C)** Finely dissected LN with surrounding fat removed.

##### Fine Lymph Node Dissection and Fixation

1. Move the Petri dish containing PBS and lymph node tissues to the stereoscope and place a Kimwipe next to the microscope for unwanted tissues.
2. Select a tissue section and begin to tease apart the fat using forceps, keeping the tissue submerged in PBS. The lymph node will appear as a brown/yellowish bead-shaped object, and the fat will appear as small, shiny, foam-like droplets.
3. Slowly peel away the fat, wiping unwanted tissue on the Kimwipe. 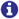 **NOTE:** Never grab the lymph node with forceps directly, as this may damage it. 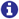 **NOTE:** It will become more difficult to manipulate the lymph node without touching it, but as fat is removed, the afferent and efferent vessels should become exposed. Manipulate the position of the lymph node by grabbing onto afferent and efferent vessels.
4. Once clean margins are achieved, move the lymph nodes to a 5 mL tube of ice-cold 1X PBS (**Figure 2C**).
5. Once all LNs are isolated, use forceps to move them into another tube with 4% paraformaldehyde (PFA) with 0.1% Triton-X overnight (approximately 18 hours) at 4°C to fix.
6. Wash lymph nodes in 1X PBS.

#### 3. Embedding and Sectioning of Lymph Nodes

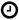 *Time: Approximately 1-2 hours, depending on the number of samples being embedded and sectioned*.

##### Embedding

1. Prepare a 6 wt% low-melt agarose solution in a glass beaker and heat in a microwave for about one minute to dissolve the agarose. 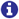 **NOTE:** The solution may boil over quickly, so watch it while in the microwave. If the bubbles reach near the top of the beaker, stop the microwave and restart once the bubbles have dissipated.
2. Place fixed lymph nodes into 10×10×5 mm vinyl cryo-molds (TissueTek, Sakura, Torrance, CA). 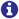 **NOTE:** If possible, place organs concave-side down (**Figure 3A**). This orientation facilitates uniform slicing of the organ and reduces the risk of the organ being pulled from the agarose during slicing. However, the anatomy of each lymph node is different and might prevent this orientation. (**Figure 3A,C**).
3. Remove PBS droplets from the LN by dabbing with a Kimwipe. 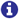 **NOTE:** The tissue should not be wetted by any left-over PBS, as this will reduce embedding strength and risk the tissue being pulled out of the agarose block during sectioning.
4. Let the agarose cool for 10 minutes at room temperature, ensuring it does not resolidify. The agarose will turn opaque as it solidifies.
5. Use a positive displacement pipette to pipette the heated agarose in a controlled manner into the mold. Place a drop of agarose in each of the four corners before quickly filling the middle, letting it create a positive meniscus slightly over the top of the well in the cryo-mold (**Figure 3C**). 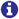 **NOTE:** This method prevents the lymph node from moving and floating to an inaccessible place in the mold. It is optimal to have the top of the node flush to the bottom of the mold so that when the block is removed from the mold, the LN is properly oriented.
6. Allow the agarose block to cool on ice. Once slightly opaque, wrap in plastic wrap to prevent drying of the samples. Samples must be sliced immediately.

**Fig. 3.**
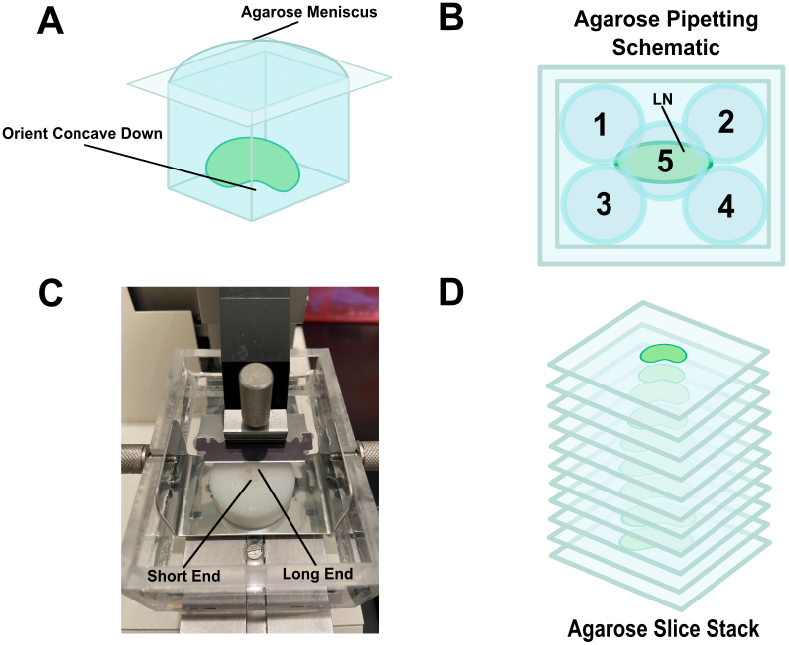
Organ embedding and mounting. **(A)** Orientation of the LN in the vinyl mold. **(B)** Order of deposition of agarose during embedding. Pipette in the four corners before flooding the center. **(C)** Orientation of the agarose block in the vibratome. **(D)** Resulting slice stack after vibratome sectioning.

##### Sectioning

1. Transport 1X PBS, a feather blade, a razor blade, super glue, and a 48-well plate filled with 0.5 mL PBS in each well on ice to a vibratome. Keep materials on ice during slicing.
2. Set up the vibratome.
3. Use the razor blade to cut the top of the agarose block flush to the cryo-mold (i.e., cut away any agarose that created the meniscus) and remove the tissue block from the mold.
4. On a hard surface, use the blade to remove the agarose block around the lymph node, leaving 2-3 mm margins. Remove a notch on the top left corner of the block to aid with proper orientation in imaging cassette. 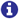 **NOTE:** This reduces the amount of time that slicing takes. The larger the agarose block, the longer it takes the blade to cut across it.
5. Put a small (size of a grain of rice) amount of superglue on the platform and set the tissue block on the platform to adhere it to the platform. Align the block so that the longer axis of the lymph node is parallel to the width of the blade (**Figure 3C**). Wait 5 minutes for the glue to dry.
6. Place the platform into the vibratome and secure it. Fill the slicing chamber with ice-cold 1X PBS.
7. Section the lymph node in 100 µm slices, using vibratome settings of 0.24 mm/s bath advance and 1800 RPM blade frequency. 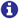 **NOTE:** Thinner slices will be more fragile, but will be quicker to stain and image.
8. Using a wet paint brush, remove the slide from the blade or PBS bath and place the tissue section with the same side up in the 48-well plate filled with PBS to keep the section hydrated. This orientation will allow users to keep track of correct tissue orientations when later reconstructing whole-organ images. 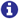 **NOTE:** Due to their fragility, slices must be manipulated with a wet paintbrush as opposed to tweezers. Wet a paintbrush in the PBS bath and use it to slide the slice off the blade, using the bristles on a corner of the agarose slice. 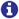 **NOTE:** Make sure to keep sections in sequential order if sectioning for 3D reconstruction (**Figure 3D**).

#### 4. Staining and Optical Clearing

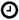 *Time: Total incubation time for staining is 96 hours (4 days)*.

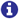 **NOTE:** The staining device should be prefilled with blocking buffer solution before adding the sample.

1. Fill staining cassette with blocking buffer.
2. Place lymph node slices orthogonally in the staining cassette in sequential order. Incubate overnight at 4°C to prevent nonspecific binding.
3. Prepare primary antibody cocktail by diluting antibodies to the following concentrations in blocking buffer:
  a. LYVE1 to 8 µg/mL
  b. MECA79 to 16 µg/mL
  c. CD31 to 16 µg/mL 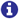 **NOTE:** Total volume needed will scale with the size of the staining cassette, but will roughly require 70 µL of reagent per 5 slices.
4. Use a pipette to remove blocking buffer and fill the cassette with the primary antibody cocktail.
5. Stain tissue slices on a rocker (roughly 30 RPM) for 48 hours at 4°C.
6. Rinse with an equal volume as antibody solution of TBST for 5 minutes on a rocker 3x at room temperature.
7. Prepare secondary antibody cocktail by diluting secondary antibodies to the following concentrations in blocking buffer:
  a. Anti-rat at 2 µg/mL (LYVE1)
  b. Anti-rabbit 10 µg/mL (MECA79)
  c. Anti-mouse 10 µg/mL (CD31)
8. Add secondary antibody solution to the well in the staining cassette, incubating on the rocker for 48 hours at 4°C. 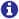 **NOTE:** Total volume needed will scale with the size of the staining cassette, but will roughly require 70 µL of reagent per 5 slices.
9. Rinse with an equal volume as antibody solution of TBST for 5 minutes on a rocker 3x at room temperature.
10. Prepare imaging cassettes for clearing by filling the wells with Focus Clear.
11. Transfer slices to the imaging cassette wells and incubate for 2 hours to optically clear the tissue slices for enhanced images on the confocal microscope. 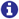 **NOTE:** Keep track of order of the slices for later 3D reconstruction 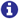 **NOTE:** There are other materials and methods for clearing tissue that may be used

#### 5. Confocal Imaging

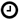 *Time: Duration of imaging can vary based on a number of factors such as number of sections, section thickness, overall size of the LN, and imaging modality used*.

1. Place another glass coverslip on top of the wells to clamp down tissue sections and prevent evaporation of the clearing solution while imaging (**Figure 4**). The weight of the coverslip is sufficient to create a lasting seal.
2. Image sections with a confocal microscope using the 488, 561, and 640 nm lasers.
  a. Objective should be chosen based on the necessary resolution for anatomical features of interest. For murine lymph nodes, long working distance 20X is recommended.
  b. Using microscope software, acquire a full-thickness tile scan of each tissue section.
    i. First, set a tile region to capture the entire surface area of the tissue. The number of tiles will scale with magnification and tissue size.
    ii. Second, set upper and lower bounds for a z-stack at each tissue face. Subdivide the z-stack into 10 sections (roughly 10 µm step-size)
3. Depending on the choice of optical clearing solution, mismatches in refractive indices may result in inaccurate measured tissue thickness in the microscope software. To correct for this, it is recommended to:
  a. Calculate scaling factor from total measured:total actual thickness ratio, and multiply the step-size by this factor, or
  b. Add fiducial marker into agar block adjacent to tissue, i.e., fluorescent microsphere with known diameter, to again calculate scaling factor from measured diameter:actual diameter ratio.

**Fig. 4.**
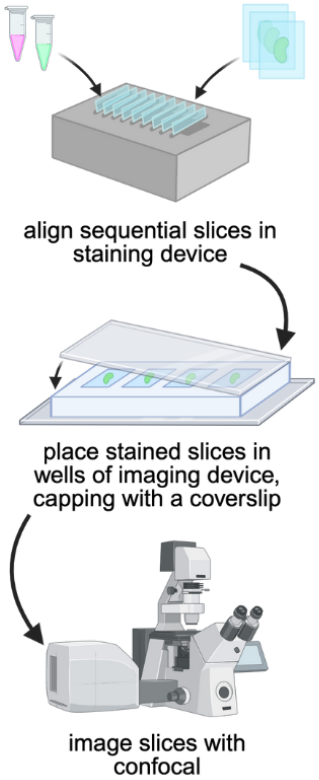
Workflow for organizing tissue slices in staining and imaging cassettes before confocal imaging. Slices are aligned orthogonally in the staining cassette for antibody staining and washing. Then they are removed and placed into the well of the imaging cassette and sandwiched with a glass coverslip to enable confocal imaging.

##### Expected Outcomes

Successful completion of this protocol will yield high-fidelity fluorescent confocal image stacks of sequential LN slices suitable for digital alignment and 3D reconstruction. Fluorescent staining reveals compartmental boundaries and vascular networks labeled by LYVE1, CD31, and MECA-79 antibodies, allowing visualization of the lobules and systemic vasculature (**Figure 5A, B**). The workflow provides reproducible slice thickness and antibody penetration, producing datasets that can be assembled in open-source platforms such as ImageJ or napari for visualization (**Figure 5C**). Similarly, the incorporation of semi-automated image registration or machine learning-based segmentation pipelines can generate 3D reconstructions for morphometric quantification, volumetric analysis, and serve as computational domains for spatial pharmacokinetic models. Because the protocol is compatible with standard confocal microscopes, it enables 3D reconstruction in laboratories lacking volumetric imaging systems, expanding access to spatial immunology and tissue-engineering studies.

**Fig. 5.**
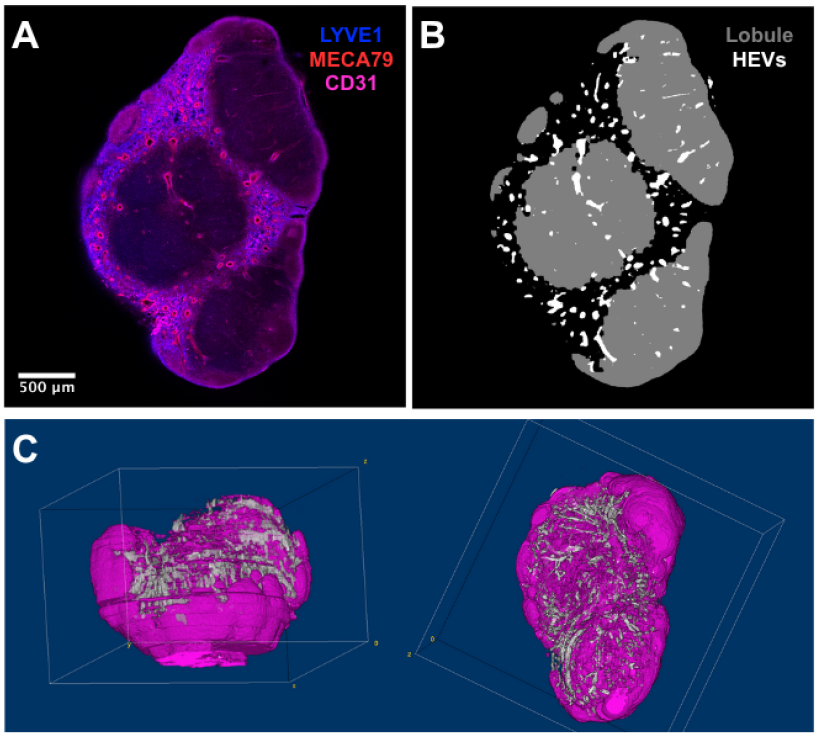
Sample image from a murine lymph node section used for 3D anatomical reconstruction. **(A)** Lymph node sections were stained and cleared using the protocol, and **(B)** segmentation masks of the lobule and high-endothelial venules (HEVs) were created from the image stack. **(C)** Segmentation masks were aligned and interpolated to generate a reconstructed 3D model (rendered using ImageJ).

## Limitations

While this workflow is designed for accessibility and reproducibility, several limitations should be considered. Manual or computer algorithm-based alignment of sequential vibratome tissue slabs introduces potential variability in Z-registration compared with automated volumetric imaging systems. Confocal imaging depth is also more limited than that of light-sheet or µCT modalities, limiting the visualization of larger organs without further sectioning. The vibratome tissue thickness will be defined by the confocal imaging depth. Additionally, because tissue morphology is defined by immunofluorescent staining, the method is limited to structures for which an antibody specific to that tissue is available.

## Troubleshooting

Troubleshooting information is compiled in Table **1**.

## DATA AND RESOURCE AVAILABILITY

Links to download SolidWorks files for staining and imaging cassettes can be found at: https://github.com/Gleghorn-Lab/Staining_Cassettes_3D_Imaging.

## ACKNOWLEDGEMENTS

The authors thank Katherine M. Nelson, Ph.D., for reviewing and commenting on drafts of the manuscript. BioRender was used in part in Figure 1 and Figure 4. This work was partly supported by grants from the National Institutes of Health U19AI158930 (JPG), R21KD142145 (JPG), and the National Science Foundation 2331440.

## AUTHOR CONTRIBUTIONS

Conceptualization (MJD, JPG), Methodology Development (MJD, JPG), Formal Analysis (MAK, MDM, MJD, JPG), Writing – Original Draft (MAK, MDM), Writing – Review & Editing (MAK, MDM, MJD, JPG), Supervision (MJD, JPG), Project Administration (JPG), Funding acquisition (JPG).

